# Evaporation and pathogenesis of levitated bacteria-laden surrogate respiratory fluid droplets: At different relative humidity and evaporation stages

**DOI:** 10.1101/2024.12.11.628080

**Authors:** Amey Nitin Agharkar, Dipasree Hajra, Kush Kumar Dewangan, Durbar Roy, Dipshikha Chakravortty, Saptarshi Basu

## Abstract

**Hypothesis:** Aerosols are the principal cause of airborne infections and respiratory diseases. Droplets ejected from the host can evaporate and form a precipitate in the air (aerosol mode), or evaporate for some time, and fall on the ground (mixed mode) or directly fall on the ground and evaporate as sessile mode. Different evaporation modes, stages of evaporation and the relative humidity (RH) conditions affect the survival and infectivity of the bacteria in the precipitate.

**Experiments:** We have investigated three droplet diameter reduction ratio-based stages of evaporation of a bacteria-laden levitated droplet at two different RH settings and evaporation modes (aerosol and mixed) mimicking real-life scenarios. The low RH condition mimics evaporation in arid regions. e.g., Delhi and the high RH conditions imitate cold areas like London. The study analyses the mass transport, micro-characterizes the samples, and investigates the survival and infectivity of bacteria in the sample.

**Findings:** The bacteria survive more in the high RH condition than in the low RH condition for all diameter reduction ratio-based stages and modes of evaporation. For the aerosol mode, at a fixed RH condition, the evaporation time plays a vital role as the bacteria in early-stage partially dried samples are more viable than the full precipitate. The evaporation rate, and the generation of reactive oxygen species (ROS) cause a remarkable difference in the viability and infectivity of the bacterial samples. Therefore, our findings report that the evaporation history of an infected droplet is an indispensable factor in determining bacterial viability and subsequent infectivity.

## 1. Introduction

The transmission of respiratory diseases via four significant modes: direct contact, indirect contact (fomites), large droplets, and fine aerosols have been well documented [1–7]. The host’s cough, sneeze, speech, etc., produce infected droplets/aerosols and moist air [8], which fall under the large droplets and aerosol transmission category of respiratory diseases. Based on droplet size and ejection speed, some droplets stay in the air for a long time till precipitation, some for a short time and settle on the surface, and some (particularly large ones) directly settle on a surface from a source [9,10]. Therefore, the events during the lifetime of a bacteria-laden droplet after ejection from an infected source is largely uncertain. Furthermore, comprehending the evaporation dynamics of tiny droplets is difficult because of their rapid evaporation [11].

Supersaturation and high surface-to-volume ratio conditions make analyzing bulk-phase aerosol jets difficult [12]. Therefore, studying a single droplet, an integral part of an aerosol jet, in an acoustic levitator answers many questions about droplet evaporation dynamics and pathogenesis. Simulation of a single droplet evaporation in an acoustic levitator is conceivable as the relative velocity-based Reynolds number between the droplet and surrounding air jet reduces to a small value very quickly [13,14].

Prevalences of bioaerosols containing pathogenic bacteria have been associated with several diseases including measles, allergies, gastrointestinal illnesses, and respiratory diseases such as influenza, asthma, and pneumonia [15–16].

Respiratory tract infections are a leading cause of morbidity and mortality around the globe [17]. 54.9% of deaths are caused by five leading pathogens-*Streptococcus pneumoniae*, *Staphylococcus aureus*, *Klebsiella pneumoniae*, *Escherichia coli*, and *Pseudomonas aeruginosa* [18]. *Klebsiella pneumoniae* is a predominant cause of antimicrobial-resistant opportunistic infections in hospitalized patients. It has been a major cause of public health threats owing to the increased prevalence of drug-resistant strains [19]. Nosocomial infections occur mainly in the respiratory tract, lungs, wound sites, and urinary tract [20]. Multivariate analyses of case reports in China showed that severe-community-acquired pneumonia (SCAP) is associated with *K. pneumoniae*, human adenovirus, human rhinovirus infection, or co-infection of respiratory syncytial virus (RSV) and haemophilus influenzae or RSV and *Staphylococcus aureus* in children and adolescents aged less than 18 years [21]. In adults, SCAP is significantly associated with infection with *K. pneumoniae*, *Pseudomonas aeruginosa*, or *Streptococcus pneumoniae*, or co-infection with *K. pneumoniae* and *Pseudomonas aeruginosa* [21]. *Klebsiella pneumoniae rhinoscleromatis*, a subspecies of *Klebsiella pneumoniae*, causes a distinctive, fatal upper respiratory disease which is often mildly contagious [22]. In our previous study [23], we had compared levitated and sessile modes of evaporation for the bacteria-laden droplet of surrogate respiratory fluid. Building on that, and motivated by the airborne modes of infections, in this work, we have rigorously studied bacteria-laden droplets of surrogate respiratory fluid in levitated mode and mixed mode (levitated and sessile) of evaporation. We study the life history of the infected droplet (refer Figure 1), starting from the source, evaporation in contact-free mode till precipitation, in contact-free mode for some duration and sessile mode for the rest (evaporation on a surface), and the droplet directly settled on fomites as sessile droplets. To decode the evaporation dynamics and the virulence properties of the infected droplet, we performed studies with a droplet of approximately 500 μm.

**Figure 1:**
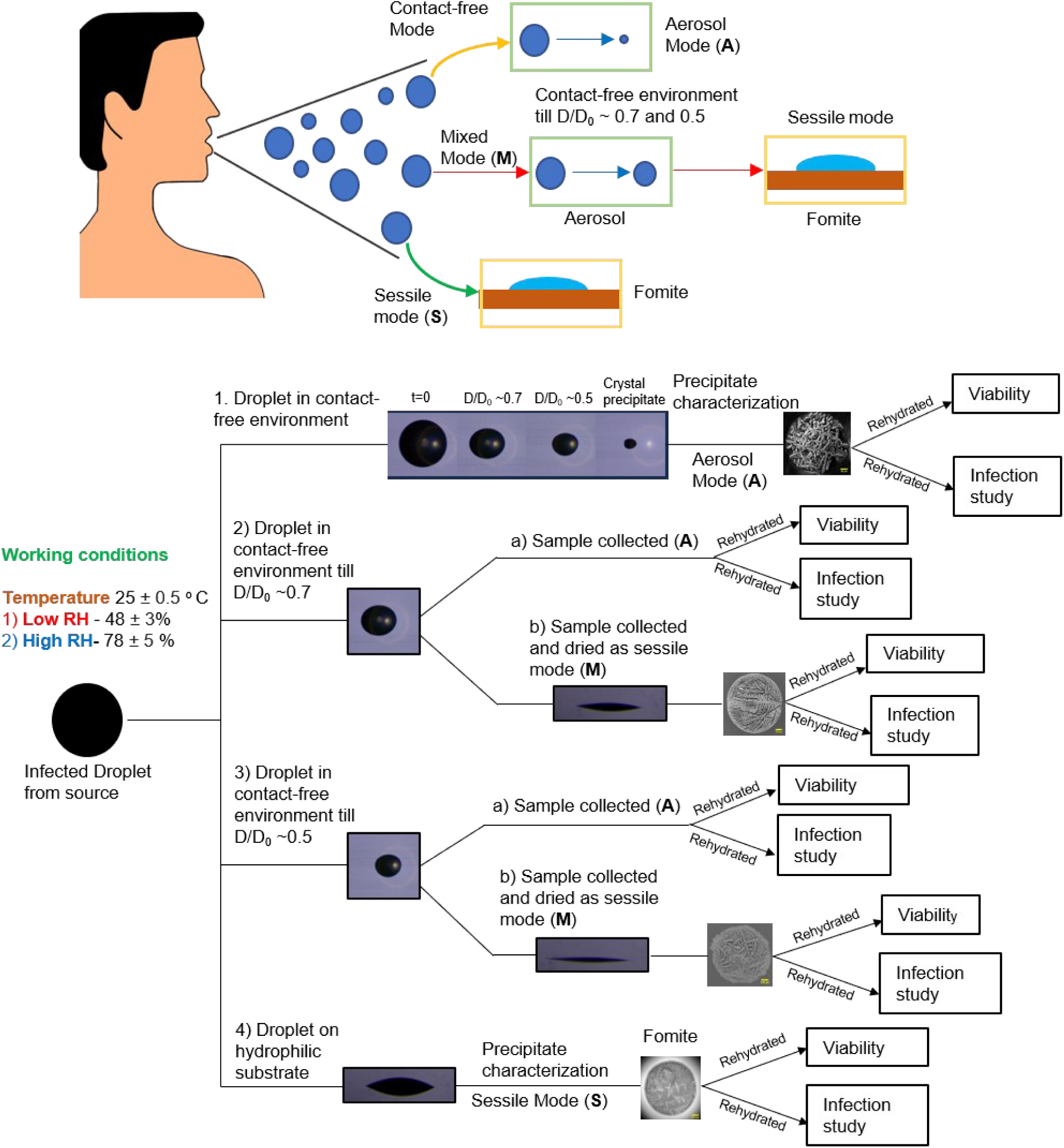
(a) Schematic of modes of infection by aerosols and droplets from an infected person (b) Infection study pathway of an infected droplet.

**Figure 2:**
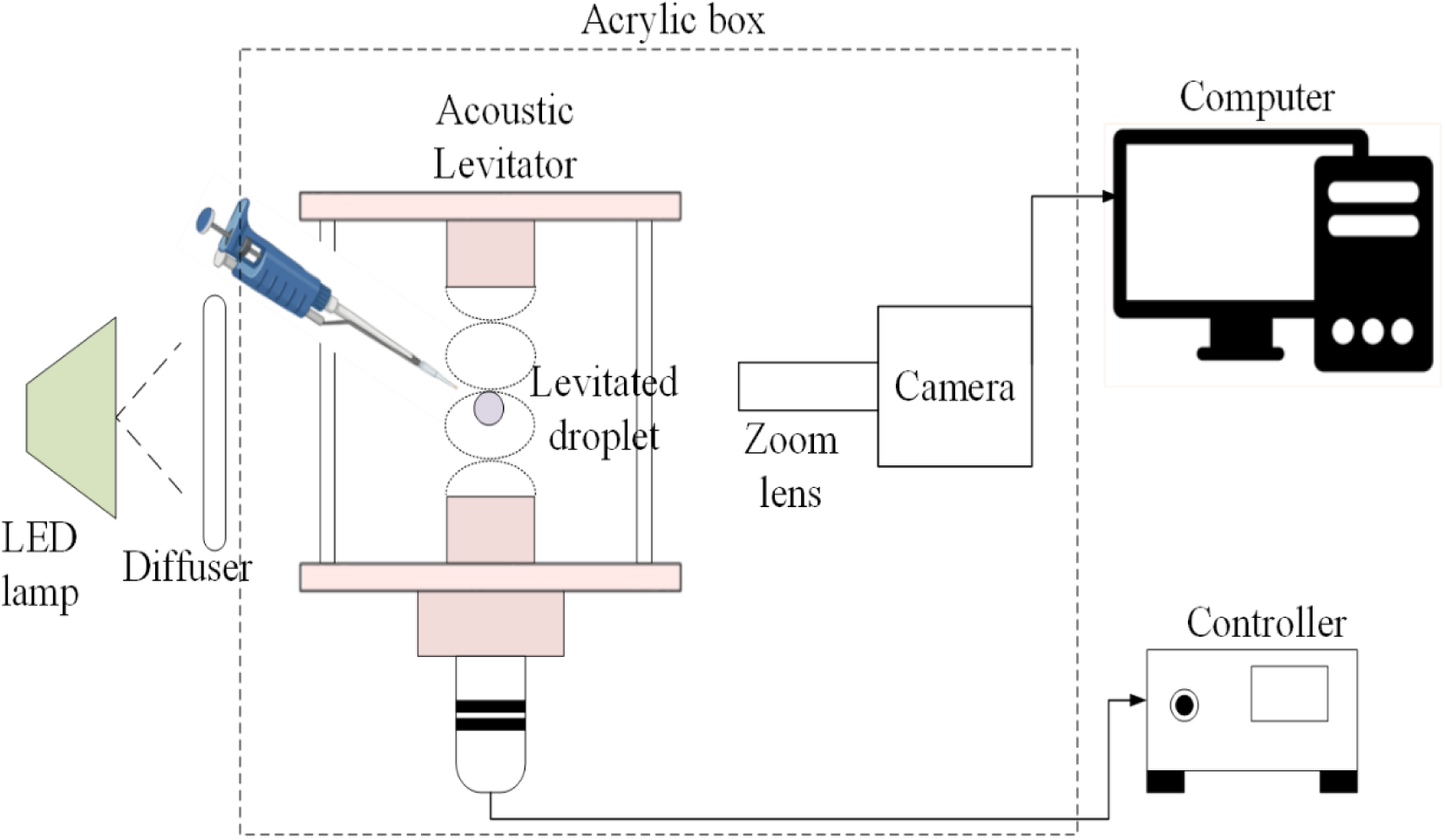
Schematic of the experimental setup for evaporation studies of Levitated droplet. The various components of the setup labelled are (1) LED lamp, (2) Diffuser plate, (3) Acoustic levitator, (4) Levitated droplet, (5) Zoom lens, (6) Camera. (7) Computer (8) Controller, and (9) Acrylic box.

Figure 1a depicts the modes of respiratory disease transmissions from an infected person. We have adapted the infected droplet evaporation occurring in the contact-free environment as aerosol mode (A), infected droplet evaporation partially in the contact-free environment, and rest as sessile droplet as mixed mode (M), and droplet settling on the surface directly and evaporating as sessile droplet as sessile mode (S). We have compared levitated droplet samples (refer Figure 1b Case 1,2a,3a) for two different RH conditions which were collected at three stages, D/D_0_ ∼ 0.7, 0.5, and precipitate (crystal), where D is the instantaneous diameter of the droplet during the evaporation process, and D_0_ is the initial droplet diameter. For the mixed evaporation mode (refer Figure 1b Case 2b,3b), the samples were collected at D/D_0_ ∼ 0.7 and 0.5 and were desiccated as sessile droplets. The evaporation studies and the micro-characterization of the samples is performed to elucidate the physics of evaporation and transport processes and the effect of evaporative stress on bacteria.

## 2. Experimental Methods

### 2.1 Sample Preparation

#### 2.1.1 Surrogate respiratory fluid preparation

The surrogate respiratory fluid (SRF) was prepared by adding 0.9% by wt. of NaCl, 0.3% by wt. of gastric mucin (Type III, Sigma Aldrich), and 0.05% wt. of di-palmitoyl-phosphatidyl-choline (DPPC) (Avanti Polar Lipids) in deionized water [24]. The final formulation was sonicated (TRANS-O-SONIC Sonicator) for 15 minutes and centrifuged (REMI R-8C) at 5000 rpm for 15 minutes [25].

#### 2.1.2 Bacterial sample preparation

Overnight grown stationary phase culture of *Klebsiella pneumoniae* MH1698 were taken, and their absorbance was measured as optical density at 600 nm using spectroscopic technique OD600nm. 10^9^ CFU (Colony Forming Units) of the bacterial culture were pelleted down at 6000 rpm for 6 minutes. The pellets were washed twice with phosphate-buffered saline (PBS) pH 7.0 and finally resuspended in 500 µL surrogate respiratory fluid (SRF).

#### 2.1.3 Sample collection procedures

The initial diameter (D_0_) of the droplet for all the cases is maintained as 660 ± 40 μm in the acoustic levitator. The aerosol mode samples (A) are collected on a microscopic cover glass (BLUE STAR, square 22 mm) at precipitate stage (crystal), D/D_0_ ∼ 0.7 and D/D_0_ ∼ 0.5 stages. The mixed mode samples (M) are collected on a microscopic cover glass at D/D_0_ ∼ 0.7 and D/D_0_ ∼ 0.5 stages, and were allowed to desiccate as sessile droplet. The cover glasses were sonicated for 2-4 minutes using propan-2-ol bath and were wiped by Kimwipes (Kimberley Clark International) before usage.

### 2.2 Experimental setup

All the experiments were performed in walk-in humidity chamber (NBS-WHC-888F with outer chamber 8×8×8 ft with all accessories and imported sensors), where 2-3 person can work while maintaining the temperature and relative humidity (RH). An ambient temperature of 25 °C ± 0.5 °C and two RH conditions were maintained: Low RH condition of 48 ± 3% and High RH condition of 78 ± 5%. Temperature and RH was measured and validated using TSP-01 sensor, Thorlabs. Figure 1 shows the experimental setup. The droplet is placed below the node in the ultrasonic levitator (tec5) using a syringe (DISPO VAN insulin syringe 31G, 0.25 mm ×6 mm of 1 mL). The lifetime of the levitated droplet is captured using a commercial DSLR camera (Nikon D5600) fitted with a Navitar zoom lens assembly (2 X lens×4.5 X tube) at 30 frames/second and a spatial resolution of 1 pixel/μm. The acoustic levitator experiments are carried inside an acrylic box to avoid convection currents of the walk-in humidity chamber. A diffuser plate is inserted between a NILA ZAILA LED light source (40 W) and the acoustic levitator for uniform illumination. The effective diameter was estimated as 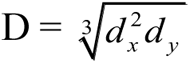, where d_x_ is the major diameter of the droplet and d_y_ is the minor diameter of the droplet. The temporal evolution of the droplet diameter during evaporation was extracted using the “Analyse particles” plugin of Image J (open-source image processing software).

### 2.3 Precipitate characterization and analysis

#### 2.3.1 Confocal microscopy sample preparation & methods

Confocal samples were prepared using the overnight grown stationary phase culture of *Klebsiella pneumoniae* MH1698. The bacteria were pelleted and washed twice using PBS and resuspended in PBS. The bacterial concentration was maintained at 10^9^ CFU/mL using OD 600. *Klebsiella pneumoniae* was stained with FM4-64 dye (1µg/mL) for 15-20 min at 37°C protected from light before resuspension in 500 µL SRF fluid. Zeiss LSM 880 NLO upright multi-photon confocal microscope was used to image the levitated sample precipitates at 10X magnification. Maximum intensity projection images were generated using the ZEN Black software (Carl Zeiss) to study the bacterial deposition in the precipitates.

#### 2.3.2 Scanning Electron Microscopy (SEM) of the samples

Similar protocol was followed for the SEM sample preparation and the bacterial concentration was maintained as 10^9^ CFU/mL. The base fluids used were SRF, 0.9% wt. NaCl in Deionized (DI) water, 0.3% wt. Mucin in DI water and bacteria in only DI water. The samples were collected from the acoustic levitator at D/D_0_ ∼ 0.7, 0.5 and precipitate stages. The samples were collected on microscopic cover glass (BLUE STAR, square 22 mm) which was coated with black carbon tape for fixing the precipitate. JEOL-SEM IT 300 Microscopy facility was used for Back scattered electrons (BSE) imaging. The device operates with a high probe current of 30 nA, acceleration voltage of 15 kV with a working distance of 12 mm.

#### 2.3.2 In vitro bacterial viability assessment

The collected levitated samples at the varied stages and modes of evaporation were reconstituted in 10 µL of sterile, autoclaved PBS. 990 µL of sterile PBS was added to the 10 µL retrieved droplet and 100 µL of it was subjected to plating on LB agar plates. Similarly, 100 µL of the bacterial sample (10^9^ CFU units in 500 µL SRF) were plated onto the respective plates at dilution of 10^-7^ and 10^-8^ which served as the pre-inoculum for normalization by taking into account the volume of the droplet (approximately 0.15 µL). 16 hr post plating and incubation at 37 °C incubator, the bacterial count was enumerated. The survival percentage was calculated as:

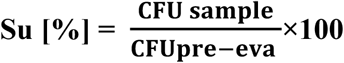

**CFU sample is the CFU of the reconstituted droplet sample.**

**CFU pre-eva is the CFU of the droplet equivalent bacterial solution sample prior to evaporation**

#### 2.3.3 Cell Culture and Infection studies

A549 lung epithelial cells were used for the infection studies. The cells were cultured in DMEM (Lonza) containing 10% Fetal Bovine Serum (Gibco) at 37 °C in a humified incubator with 5% CO_2._12 hr -14 hr prior to each experiment, cells were seeded into 24 well at a confluency of 60%. The bacteria reconstituted from the samples were used for infecting the cells. The infected cells were incubated in 5% CO_2_ incubator for 25 min at 37 °C. Post-incubation, the infected cells were washed with sterile PBS and were incubated at 37 °C in a 5% CO_2_ incubator for 1 hr in the presence of 100 µg/mL gentamicin containing DMEM media. Post 1hr incubation, cells were further incubated with 25 µg/mL gentamicin containing DMEM media at 37°C in the 5% CO_2_ incubator after a PBS washing step. At designated time points post-infection, cells were lysed with 0.1% Triton-X-100 in PBS. The lysates were serially diluted and plated on LB agar plates. The CFU at the later time points of infection (16 hr) was divided by the corresponding CFU at the initial time points of infection (2 hr) to calculate the fold proliferation.

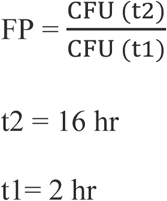

#### 2.3.4 Flow Cytometry

The samples were reconstituted in 1X PBS. The reconstituted samples were centrifuged at 6000 rpm for 6 minutes to obtain the bacterial pellet. The pellets were resuspended in the DCFDA (20µM) staining solution and incubated for 15 minutes in dark at 37 °C. Following incubation, centrifugation was performed at 6000 rpm for 6 minutes to obtain the stained bacterial pellet. The bacterial pellets were washed in 1X PBS and eventually resuspended in 100 µL of 1X PBS for FACS recording. The FACS recording was performed in BDFacsVerse instrument and analysis was performed in BDFacsSuite software.

#### 2.3.5 Statistical analysis for viability, infection and cytometry studies

Data were analyzed, and graphs were plotted using the GraphPad Prism 8 software (San Diego, CA). Statistical significance was determined by Student’s t-test to obtain p values. Adjusted p-values below 0.05 are considered statistically significant. The results are expressed as mean ±SEM (Standard Error of Mean).

## 3. Results and Discussion

### 3.1 Evaporation dynamics of the levitated droplet

Our previous work of bacteria-laden levitated versus sessile droplets show that the recipient ingesting levitated precipitate is more virulent than the sessile precipitate. Building on that, to do in-depth research on bacteria-laden droplets from an infected host, we focused on the different stages of levitated droplet evaporation (aerosol mode) at two RH conditions. We extended the study to mixed evaporation mode (levitated and sessile).

The droplets’ volume reduction rate during evaporation is the consequence of the evaporation rate [23]. As we know, the evaporation rate depends on physical parameters such as droplet temperature, ambient temperature, relative humidity (RH), droplet size, vapor pressure, mass diffusivity, thermal diffusivity, surface geometry, and solute properties; we have experimented with two different RH conditions, maintaining the ambient temperature and initial solute concentration constant. To decipher the effect of evaporation time on bacterial viability and infectivity with respect to initial droplet, the droplet sample was collected at three stages during the droplet evaporation: D/D_0_ ∼ 0.7, 0.5 and precipitate. Also, the D/D_0_ ∼ 0.7 and 0.5 samples were collected during evaporation and were allowed to dry as sessile droplets, mimicking a mixed mode of evaporation (see Methods). The droplet length scale is an important parameter governing the evaporation process occurring either in diffusion regime or reaction regime. Since the initial droplet diameter in our study is of the order of O (10^2^) μm and the final precipitates of order of O (10^1^) μm, our experimental study follows the diffusion limited regime of droplet evaporation.

Figure 3 represents the volume ratios V/V_0_ (V is the instantaneous volume and V_0_ is the initial droplet volume) with respect to time in seconds of the evaporating droplet in the acoustic levitator. Figure 3 represents the two fluids’ low RH and high RH experimental data, respectively. The green color plot and the blue plot are for KP in the SRF sample (bacterial sample), abbreviated as KP + SRF for low RH and high RH cases, respectively. The brown color plot and red plot describe only SRF (Base fluid) droplet regression abbreviated as SRF for low RH and high RH cases, respectively. The insets in Figure 3 show the droplet at different diameter reduction ratio-based stages. Inset (i) depicts the initial droplet, (ii) shows the state of the droplet at D/D_0_ ∼ 0.7, (iii) of the droplet at D/D_0_ ∼ 0.5, and (iv) represents the droplet after precipitation (crystallization). The High RH droplet requires more time to evaporate around t ∼ 1600-1750 s and the low RH samples evaporate faster around t ∼ 800-900 s. The evaporation process occurs in two stages: fast initial evaporation and slow evaporation towards the end. The droplet size becomes stagnant as the evaporation rate decreases towards the end of evaporation because of the increased solutal concentration at the interface [26]. Subsequently, the evaporation occurs from the interstices of the solute skin. On reaching the efflorescence limit, the droplet crystalizes very fast.

**Figure 3:**
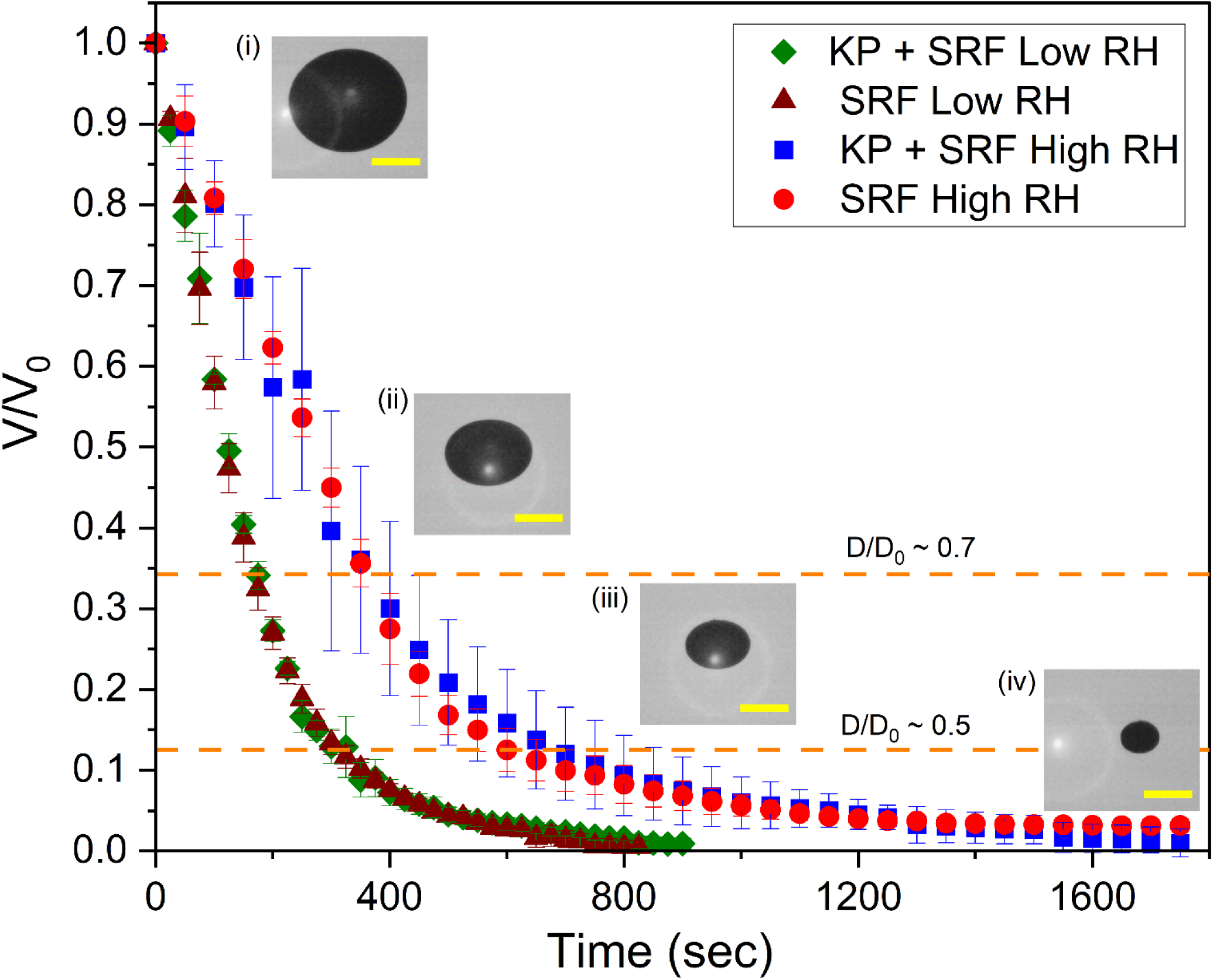
Volume ratio V/V_0_ variation as a function of time in seconds for the levitated drop evaporation. Here green color describes KP + SRF droplet at low RH, brown color describes SRF droplet at low RH, blue color describes KP + SRF droplet at high RH and red color describes SRF droplet at high RH. Insets show the droplet side view images at various stages. (i) Initial Droplet (ii) Droplet at D/D_0_ ∼ 0.7 (iii) Droplet at D/D_0_ ∼ 0.5 (iv) Droplet after precipitation (crystallization). All Scale bars represent 200 μm. Orange color dashed lines represents D/D_0_ ∼ 0.7 and 0.5 stages on V/V_0_ graph respectively.

The experimental average evaporation rates for low RH droplet case and high RH droplet case are: 1.77×10^-10^ kg/s and 9.77×10^-11^ kg/s respectively. Experimentally the evaporation rate of low RH levitated droplet is one order higher than the high RH droplet. The RH governed evaporation rate which in turn affects the viability and infectivity of bacteria in a levitated droplet. (refer Section 3.3)

For the levitated droplet configuration, the evaporative flux is uniform and the acoustic streaming effects generates the majority of the toroidal flow field within the drop [27,28]. The toroidal flow velocities inside a levitated drop are of the order of mm/s-cm/s [29]. A study showing the transition of internal and external flows of 50 wt. % ethanol-based water droplet with evaporation confirmed that the evaporation rate affects the droplet’s internal flow [30]. Therefore, the flow in acoustically levitated low RH droplet is more intense than the high RH droplet because of high evaporation rate. The concentration at the receding edge is higher in low RH case than high RH case due to the transport of solute towards the interface, affecting the packing and survival of bacteria.

The stress experienced by the bacteria due to evaporation of water in the bacteria-laden droplet is defined as the evaporative stress. The evaporation rate of the droplet is a marker of evaporative stress. Furthermore, the stress experienced by the bacteria inside the droplet due to the acoustic pressure on the droplet is termed as acoustic stress. Towards the end of the evaporation in general levitation experiments, the buckling of the droplet occurs causing deformations in the final precipitates [31,32]; therefore, the stress experienced by the bacteria due to buckling of the precipitate is known as buckling stress. In our case, the buckling pressure on the bacteria is three orders lower than the acoustic pressure; hence it has negligible effect (refer Section 3.2 for scaling analysis). Therefore, the evaporative stress and acoustic stress play an important role on the bacterial survival.

Important parameters for the survival of bacteria in samples are: Firstly, the evaporation time plays a crucial role. For the aerosol mode droplet at the constant RH condition, the bacterial survival in D/D_0_ ∼ 0.7 sample is more than the D/D_0_ ∼ 0.5 sample and final precipitate (fully desiccated) sample. More the droplet evaporation, more the water loss (mass loss); hence, the bacteria experience more evaporative stress. The bacterial inactivation occurs due to osmotic pressure from increased salt concentration in the droplet due to water loss during evaporation process [33]. Therefore, the evaporation time plays a significant role in the diameter reduction ratio-based stage-wise droplet evaporation.

Secondly, the RH governed evaporation rate defines the evaporation time for the droplet evaporation. The low RH droplet evaporates faster while the high RH droplet takes a longer time. Longer the evaporation time, lesser is the evaporative stress; the bacteria in essence get enough time to absorb nutrients from the SRF base fluid and form aggregate networks.

Thirdly, for the mixed mode of evaporation droplet, the substrate geometry plays a vital role. After being in the contact-free environment (levitated) starting from the initial diameter (D_0_), the droplet is collected at D/D_0_ ∼ 0.7 and 0.5. Subsequently, the evaporation proceeds as a sessile droplet. From our previous study, we know that the evaporative flux at the contact-line of the sessile droplet is maximum because of the coffee-ring effect. The bacteria in the mixed-mode samples for a fixed RH condition, experiences more evaporative stress than the corresponding aerosol mode droplet sample, hence the survivability is affected. The bacteria in the mixed mode samples have lower survivability than the aerosol mode sample (contact free throughout the evaporation lifetime). For the pure sessile case, the droplet is into the sessile configuration of evaporation since its inception, therefore it would have lower survivability and infectivity than aerosol mode precipitate at a fixed RH condition.

The mode of evaporation, evaporation geometry, relative humidity, stage of the droplet sample collection etc. are the major factors affecting the evaporation rate and consequently the survivability of bacteria in samples.

### 3.2 Micro-characterization of dried samples

Figure 4 shows the maximum intensity confocal microscopy images of the dried samples of bacteria-laden droplet samples in mixed mode of evaporation, collected at D/D_0_ ∼0.7 and 0.5 for both the RH conditions. Figure 4a represents mixed mode precipitates at D/D_0_ ∼ 0.7 for low and high RH conditions. Similarly, figure 4b depicts mixed mode precipitates at D/D_0_ ∼ 0.5 for low and high RH conditions. The plots in figure 4a and 4b shows the variation of the fluorescence intensity in arbitrary units versus spatial location ratio (x/D, where x is distance from one edge of the droplet precipitate to another and D is the diameter of the droplet precipitate) for a band of 100 μm in width. From the figures and the plot, we can observe the accumulation of bacteria at the edge of the droplet precipitate for both the stages of low RH case, while for both the stages of high RH case, the distribution is diffused all over the droplet. After the droplet is collected at D/D_0_ ∼ 0.7 and 0.5 stages, the droplet evaporates as a sessile droplet. During the sessile droplet evaporation, the flow is dominated by capillary forces (coffee ring effect) towards the edge of the droplet [34]. The mucin present in the SRF absorbs water vapor, and it is more prone to absorb at the high RH condition than the low RH condition, resulting in a diffused and broader edge distribution in the high RH precipitate [35]. Not only the evaporation rate but also the pathogen distribution in the precipitate influences its viability, as the deposition locations are important for differential desiccation [36,37]. Therefore, the mixed mode viability results depict higher survival of high RH samples than the corresponding low RH samples. (refer Figure 8a).

**Figure 4:**
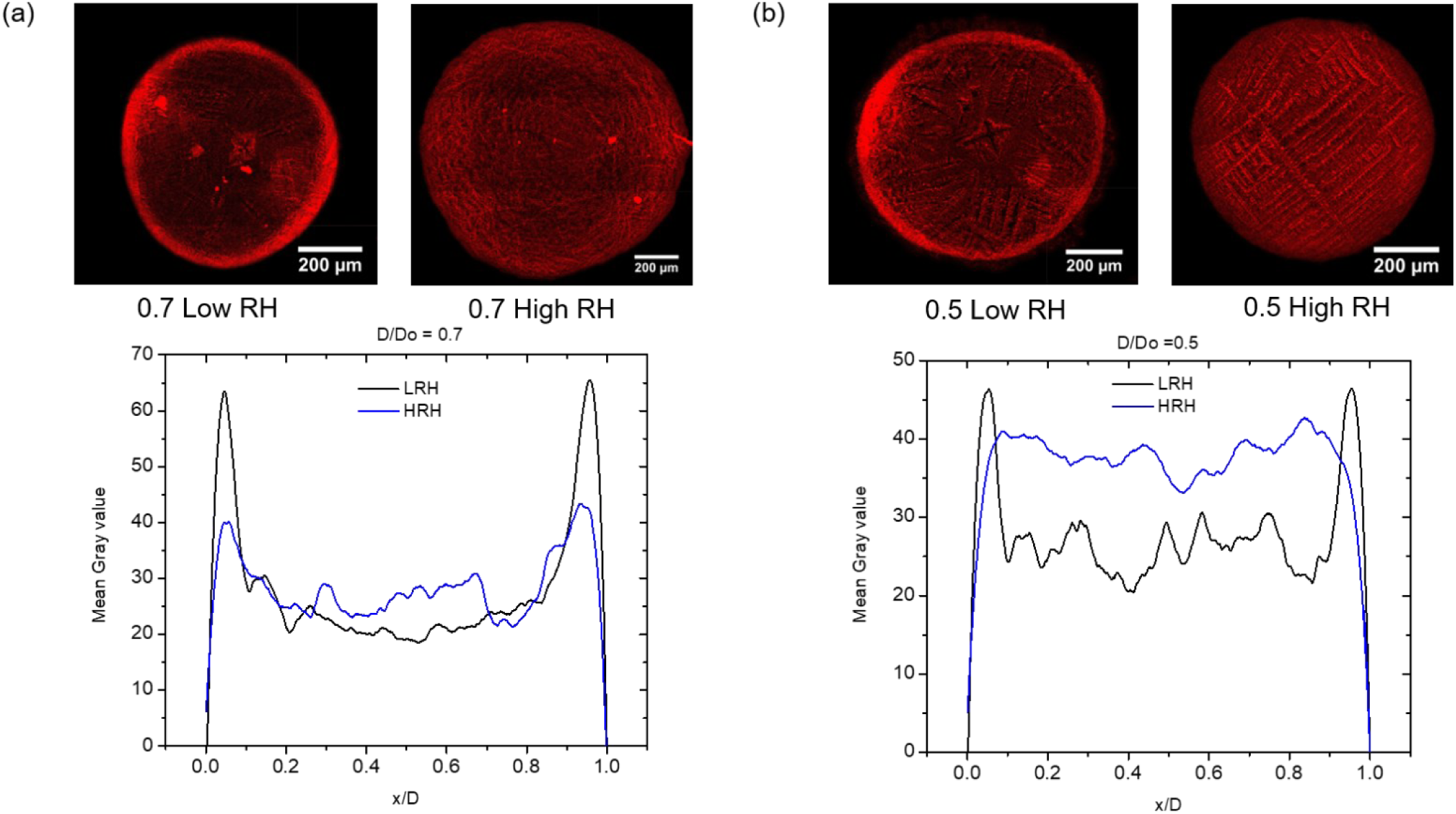
Confocal microscopy of mixed mode evaporation dried samples with intensity scales: (a) D/D_0_ ∼ 0.7 mixed samples for low and high RH (Case 2b) (b) D/D_0_ ∼ 0.5 mixed samples for low and high RH (Case 3b).

Figure 5 shows the SEM images of the mixed mode samples for both the RH conditions and both the stages of D/D_0_ ∼ 0.7 and 0.5. Inset (i) depicts the centre of the precipitate and Inset (ii) depicts the edge of the precipitate for all the cases. Multiple thin equiaxed dendritic structures with fine branches are formed in the low RH precipitates (refer Figure 5 a, c), while in high RH precipitates the dendrites are thicker and originate from a single equiaxed dendritic nucleate. (refer Figure 5 b, d) [35]. The NaCl crystal nucleation occurs faster in low RH case due to higher evaporation rate, therefore small dendrites are formed. For the high RH case, the nucleation starts from the centre as a single dendrite, since the evaporation rate is low [35].

**Figure 5:**
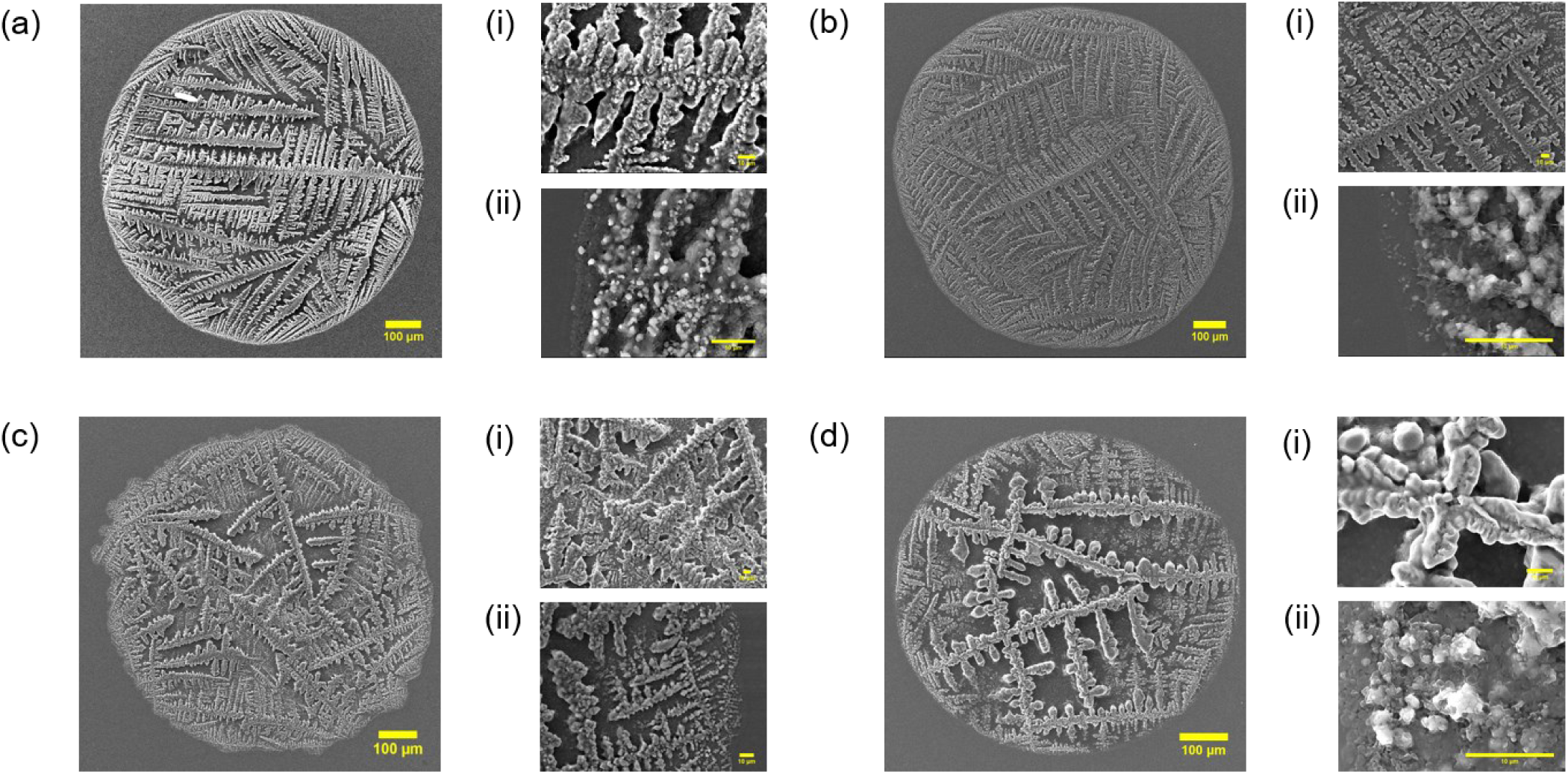
(a) SEM image of mixed mode Low RH KP + SRF precipitate collected as levitated at D/D_0_ ∼ 0.7, then sessile mode (i) SEM at centre (ii) SEM at edge (b) SEM image of mixed mode High RH KP + SRF precipitate collected as levitated at D/D_0_ ∼ 0.7, then sessile mode (i) SEM at centre (ii) SEM at edge (c) SEM image of mixed mode Low RH KP + SRF precipitate collected as levitated at D/D_0_ ∼ 0.5, then sessile mode (i) SEM at centre (ii) SEM at edge (d) SEM image of mixed mode High RH KP + SRF precipitate collected as levitated at D/D_0_ ∼ 0.5, then sessile mode (i) SEM at centre (ii) SEM at edge

Figure 6 depicts the SEM image, side view image and the confocal image of the levitated precipitate (aerosol mode). The low RH precipitate side view image (refer Figure 6 a (iii)) shows precipitate with jagged edges, while high RH precipitate (refer Figure 6 b (iii)) seems to be like a cocoon shaped structure. The evaporation rate plays a major role in governing the internal flow and the arrangement of the non-volatile solutes in the precipitate. The mass Peclet number analysis explains the phenomena of the bacterial distribution in the levitated droplet [14,31].

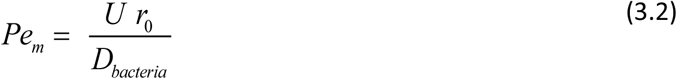

**Figure 6:**
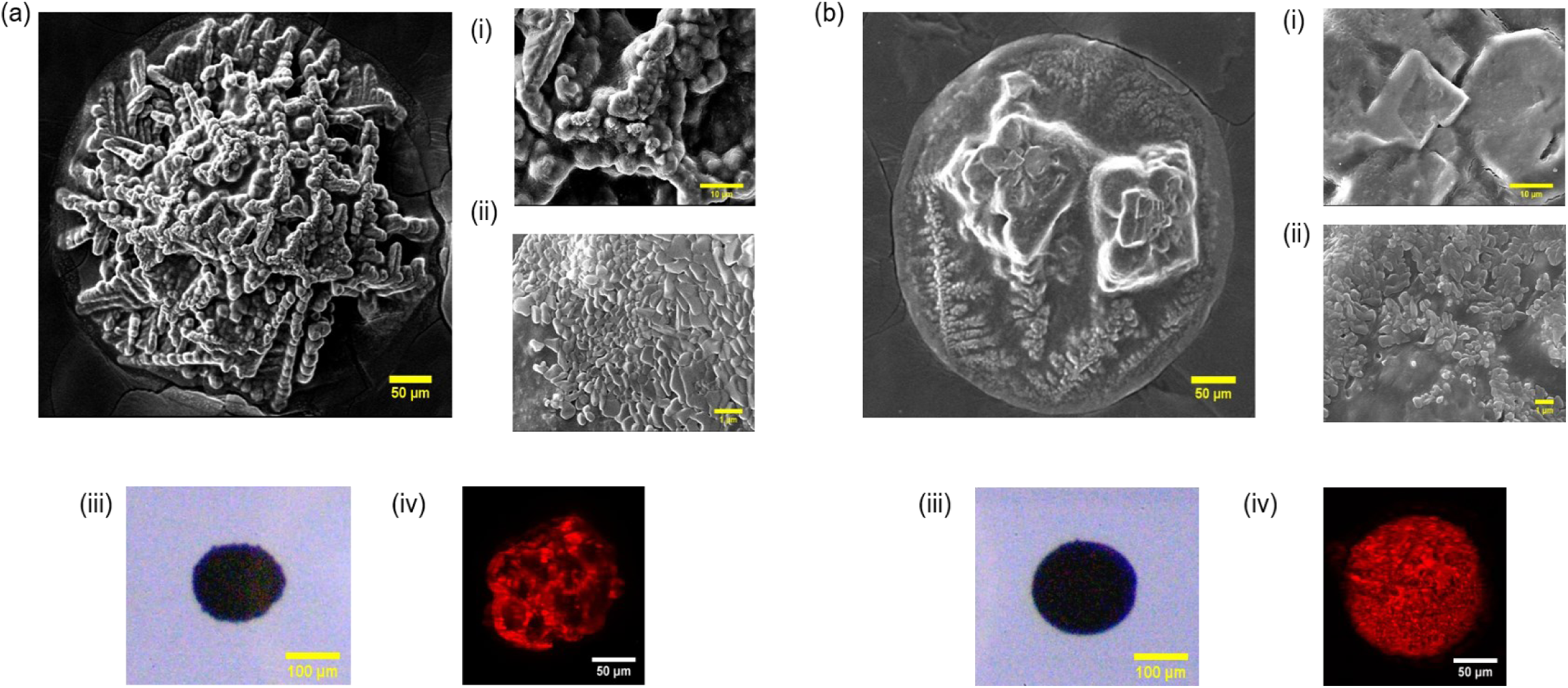
(a) SEM image of Low RH KP + SRF precipitate (i) SEM at centre (ii) SEM at edge (iii) Side view image of Low RH KP + SRF precipitate in acoustic levitator (iv) Confocal image at maximum intensity projection of Low RH KP + SRF precipitate. (b) SEM image of High RH KP + SRF precipitate (i) SEM at centre (ii) SEM at edge (iii) Side view image of High RH KP + SRF precipitate in acoustic levitator (iv) Confocal image at maximum intensity projection of High RH KP + SRF precipitate.

Where U is the rate of droplet diameter reduction (U ∼ 0.44 μm/s for low RH and U∼ 0.32 μm/s for high RH), r_0_ = 330 μm is the initial droplet radius, and D_bacteria_ is the mass diffusivity of bacteria in SRF (assuming bacterial transport similar to particle transport), which is calculated using Stokes-Einstein equation:

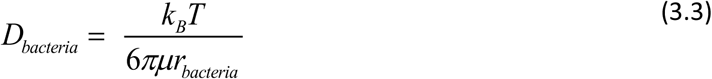

Where k_B_ is the Boltzmann constant, T = 298 K is the ambient temperature, μ = 1.23×10^-3^ Pas is the dynamic viscosity of SRF, r_bacteria_ = 0.5 μm is the assumed bacterial radius.

We get D_bacteria_ = 3.55×10^-13^ m ^2^/s, leading to Pe_m_ = 4.15×10^2^ for the low RH case and Pe_m_ = 2.98×10^2^ for high RH case. Therefore, Pe_m_ ∼ O (10^2^). For both the low and high RH cases, Pe_m_ >> 1 implies bacterial accumulation near receding interface of the droplet and there is no diffusion in the droplet while evaporating. The mass Peclet number at low RH condition is higher than high RH condition, implying higher concentration of the non-volatile solutes at the receding edge. This phenomenon was confirmed by the Energy-dispersive spectroscopy (EDS) reports for both the samples (refer Supplementary Figures 2, 3). The high RH precipitate has more homogeneous distribution of Carbon (C), Oxygen (O), Sodium (Na) and Chlorine (Cl) at the centre of the precipitate and the edge of the precipitate. While for the low RH precipitate the centre has dominant presence of Sodium (Na) and Chlorine (Cl), confirming heterogeneous distribution. The florescent images of confocal microscopy (refer Figure 6 a (iv) and 6 b (iv)) displays heterogeneity in bacterial distribution in low RH precipitate than the high RH precipitate. The rapid nucleation and transport of Na^+^ and Cl^-^ ions in the low RH precipitate governed by the high evaporation rate causes decrease in bacterial survivability compared to high the RH precipitate. The increase of solute concentration leads to the inactivation of bacteria due to increase in osmotic pressure [33].

Towards the end of the evaporation, when the droplet size gets fixed, the evaporation happens through the pores of the solutal skin. This can lead to the buckling of the droplet precipitate and form precipitate shapes according to the evaporation rate [32]. Therefore, we evaluated the bucking pressure using the following equations [31,32,38,39]

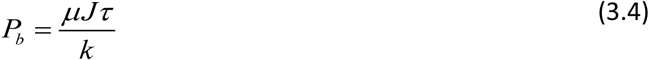

P_b_ is the capillary pressure drop obtained from the Darcy’s law, J is the solvent vaporization flux through the pores which is evaluated as 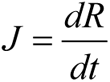 (radius reduction rate after the onset of buckling), τ is the shell thickness which is evaluated using 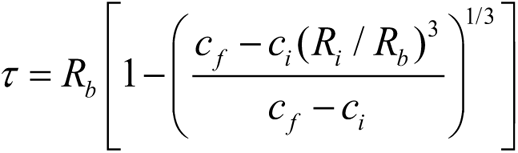 where c_i_ = 0.0143 is the initial solute concentration and c_f_ = 0.56 is the final solute concentration R_i_ = 325 µm is the initial radius of the droplet and R_b_ = 111 µm is the radius of the droplet at buckling μ = 0.00119 Pa.s is the dynamic viscosity obtained by equation 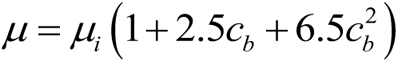, where c_B_ is *c_B_* = *c_i_* (*R_i_* / *R_b_*)^3^ and μ_i_ is the initial viscosity. Here k is the shell permeability obtained by the Kozeny-Carman equation as 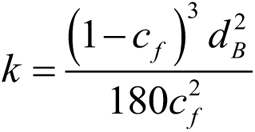 where d_B_ = 1 µm is the bacterial diameter. *J* = 2.45×10^-7^ m /s for low RH case and *J* = 1.34×10^-7^ m/s for the high RH case. The buckling pressure evaluated for both the cases is 3.8 Pa and 2.3 Pa for low RH and high RH respectively. Since the bacterial size is in µm scale, it faces less bucking stress than any nanoparticle laden droplet as shown in earlier works [31,32]. Also, the presence of mucin and DPPC inhibits buckling because of the reduction in evaporation rate and disruption in porous shell formation [32]. Buckling is caused by the combined effects of evaporation rate, solute/colloid properties and morphologies and particle diffusion. Therefore, the buckling pressure which is of order of O (10) Pa can be neglected compared to the acoustic pressure which is of order O (10^3^) Pa.

The higher evaporation rate of low RH droplet causes the precipitate to be formed with jagged edges (irregular shape). While the high RH precipitate is formed as a cocoon shaped [32].

### 3.3 The aerosol mode of evaporation at high relative humidity facilitates increased bacterial survival, exhibits increased intracellular proliferation in infected lung epithelial A549 cells and exhibits lower oxidative stress

To assess the survival of *Klebsiella pneumoniae* under the levitated mode of evaporation at low or high relative humidity, we performed an *in vitro* survival assay by plating the reconstituted evaporated droplets onto LB agar plates. We observed increased bacterial survival in the levitated droplets at high relative humidity (RH) compared to low RH droplets (refer Figure 7a). Moreover, larger droplets with high-diameter harbors increased bacterial number among the high RH droplets. Together, our data indicate increased bacterial survival within the levitated droplet at high RH and with high D/D_0_.

**Figure 7:**
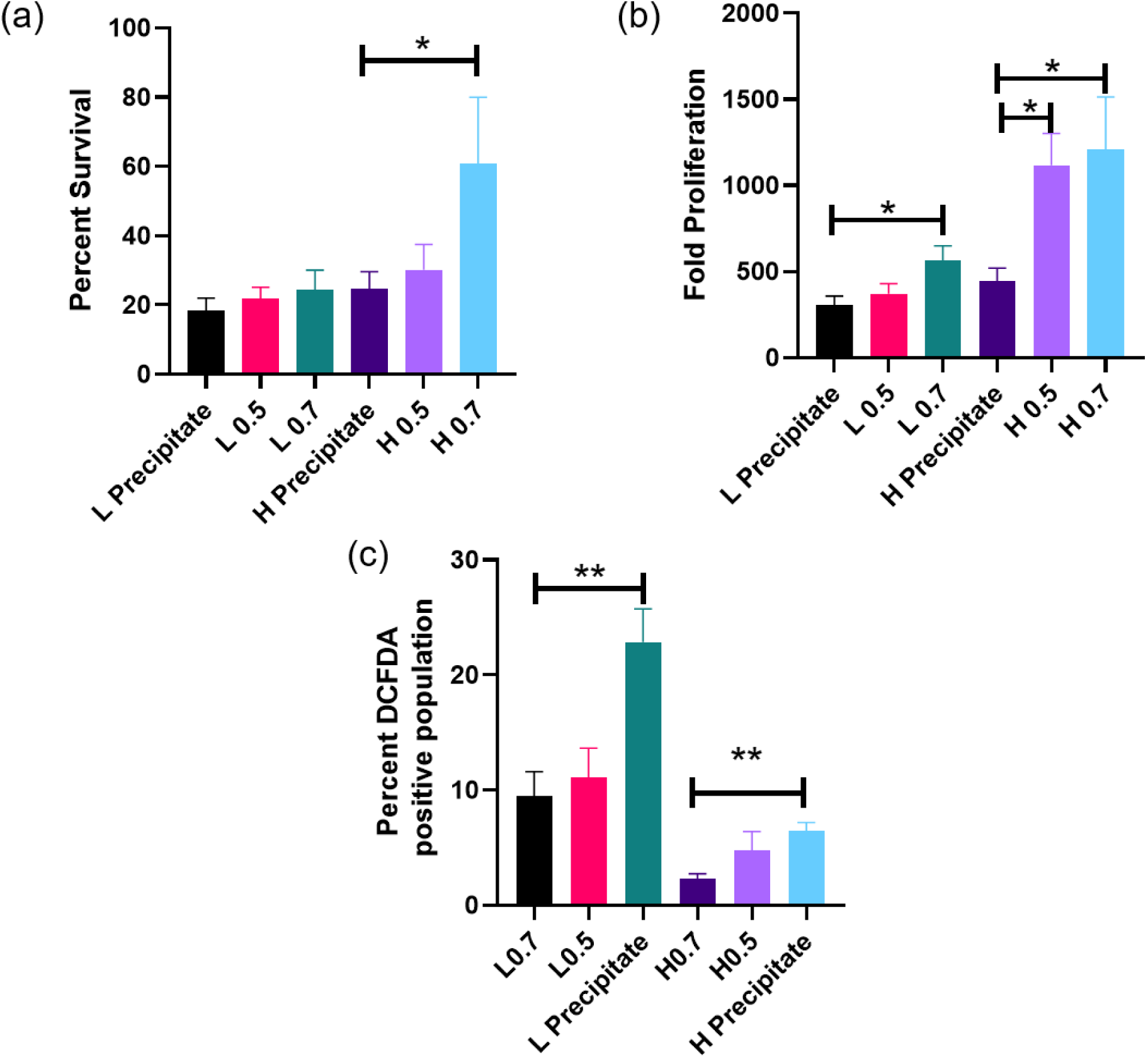
(a) *In vitro* survival of *K. pnuemoniae* in levitated samples at Low Rh (L) and High RH (H) for D/D_0_ ∼ 0.7, 0.5 and precipitate samples (Case 1). The data is representative of N=3 (biological replicates), n=3 (technical replicates). (Refer Supplementary material Section S1 for * value) (b) Intracellular proliferation of *K. pnuemoniae* for infection within lung epithelial cells A549 (N=2, n=3). (c) Flow cytometric quantitation of Reactive oxygen species (ROS) by Percent 2’,7’Dichlorofluorescein diacetate (DCFDA) positive population of *K. pnuemoniae* (N=3, n=3).

As data suggested increased survival of bacteria in the high RH levitated droplets with large D/D_0_, therefore we wanted to evaluate the bacterial replication efficiency in the lung epithelial A549 cells. The levitated bacterial droplets at high RH showed enhanced intracellular proliferation within the infected A549 cells when compared to low RH droplets (refer Figure 7b). Moreover, droplets with higher D/D_0_ ratio demonstrated higher intracellular proliferation for both high and low RH cases.

Our previous findings demonstrated lower bacterial survival in levitated under low RH conditions when compared to high RH. To investigate whether lower bacterial survival is triggered by higher production of reactive oxygen species (ROS), we estimated ROS generation using DCFDA staining via flow cytometry [40]. ROS cascade is one of the lethal stressors that leads to microbial cell death [41,42]. 9.4% and 11.05% of *K. pneumoniae* from the low RH levitated droplet with 0.7 and 0.5 D/D_0_ showed enhanced ROS production, respectively in comparison to around 22.86% of the levitated precipitate with low RH population exhibiting high ROS production (refer Figure 7c). Compared to low RH levitated droplets, the high RH droplets exhibited reduced ROS generation with only 2.31%, 4.74% and 6.44% of *K. pneumoniae* population in the high RH droplets having droplet diameter D/D_0_ range of 0.7, 0.5 and precipitate respectively (refer Figure 7c). Therefore, low ROS generation during high RH conditions facilitates higher bacterial survival both in vitro and within infected lung epithelial cells [23].

### 3.4 Levitated droplets with a mixed mode of evaporation (levitated mode of evaporation followed by sessile mode of evaporation following contact with surface) under high RH exhibit increased in vitro bacterial survival and intracellular proliferation

Further, we wanted to evaluate the effect of the mixed mode of evaporation on the *in vitro* survival of *Klebsiella pneumoniae* under high and low RH conditions. *Klebsiella pneumoniae* showed increased survival at high RH conditions (refer Figure 8a). Moreover, evaluation of intracellular proliferation of mixed sample *Klebsiella pneumoniae* within the infected A549 cells revealed enhanced intracellular proliferation under high RH conditions as opposed to low RH conditions (refer Figure 8b).

**Figure 8:**
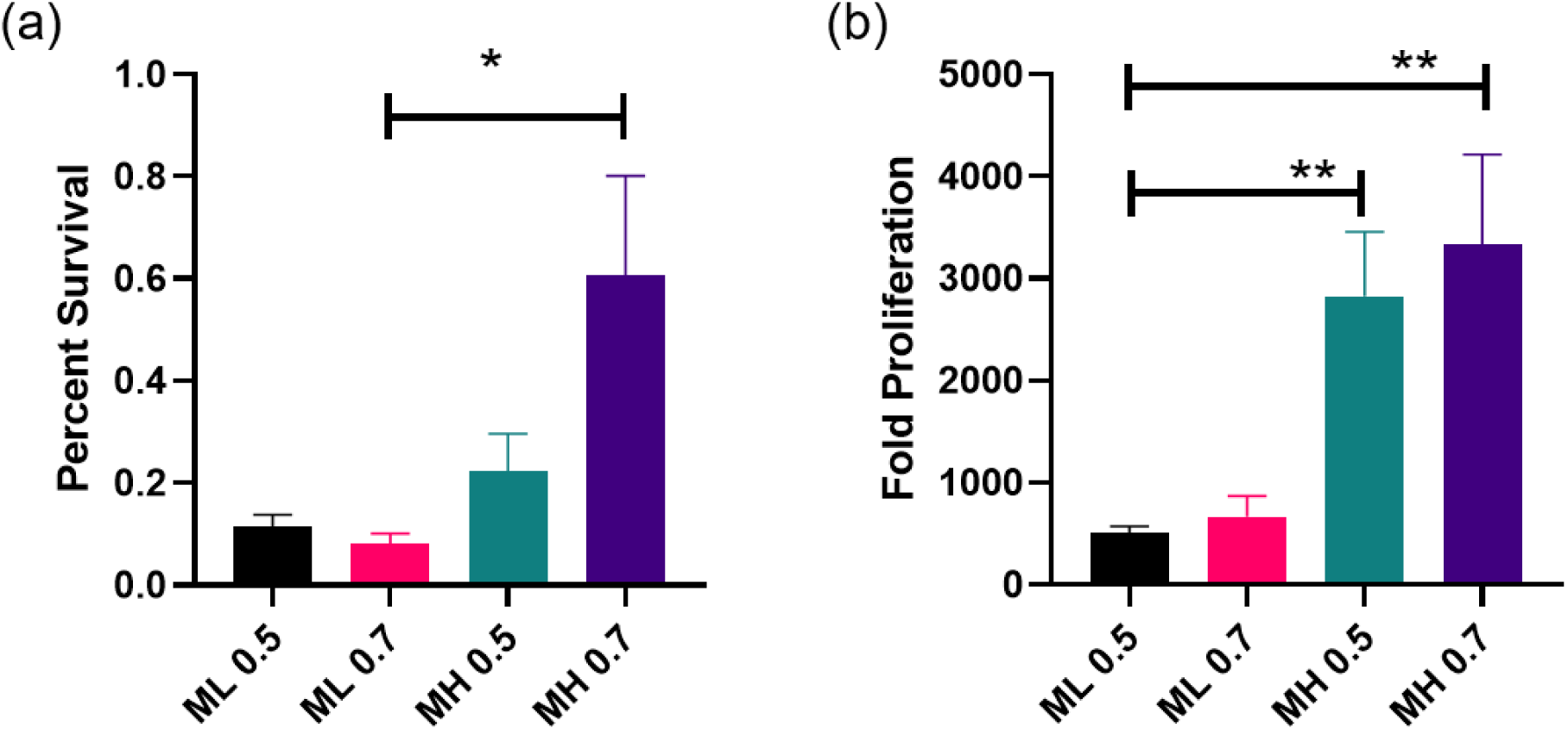
*In vitro* survival of *K. pnuemoniae* in mixed mode evaporation samples at Low RH (ML) and High RH (MH) for D/D_0_ ∼ 0.7, 0.5 and precipitate samples (Case 2b & 3b). (N=2, n=3). (b) Intracellular proliferation of *K. pnuemoniae* for infection within lung epithelial cells A549 (N=2, n=3).

## 4. Conclusions

The history of droplet evaporation is a paramount factor in deciding the distribution of bacteria in the precipitate and its viability and infectivity. The mode of droplet evaporation affects the evaporation rate and time. We show that the relative humidity-dependent evaporation rate is pivotal in bacterial survival and virulence. Whereas for the diameter reduction ratio-based stages of evaporation at a fixed RH condition, the evaporation time of the collected sample is the deciding factor for bacterial survival in sample. For the same initial diameter droplet evaporating in aerosol mode, the average mass evaporation rate for the low RH droplet is at least one order larger than the corresponding droplet at high RH. The evaporation rate affects the flow of bacteria inside droplet. In the high RH D/D_0_ ∼ 0.7 and 0.5 mixed mode precipitates, the bacterial distribution is more homogeneous than in low RH precipitates. Similarly, for the aerosol mode evaporation, the high RH precipitate bacterial distribution is more uniform than the low RH precipitate. Compared to the low RH samples, the *in vitro* comparative viability studies revealed increased survival of *Klebsiella pneumoniae* in the high relative humidity samples for an aerosol mode of evaporation. For the diameter-based stage-wise aerosol mode samples, the survivability decreases from the D/D_0_ ∼ 0.7 sample to the crystal precipitate sample for both RH conditions. Similarly, high RH aerosol droplet-laden bacteria exhibit heightened intracellular proliferation within lung epithelial cells than the low RH aerosol droplet. Furthermore, examination of the mixed mode of evaporation on the *in vitro* survival and intracellular replication of *Klebsiella pneumoniae* under high and low RH conditions depicted that high RH conditions lead to enhanced in vitro survival and intracellular fold proliferation compared to low RH.

Apart from the evaporation rate and time, ROS production contributes to the differential bacterial viability and virulence. Higher ROS production in aerosol-mode bacterial samples at low RH contributes to the lower bacterial burden compared to the high RH aerosol. Further, the longer the evaporation time of the droplet in aerosol evaporation mode, the greater the superoxide production for stage-wise evaporation cases for both RH.

In this study, we showed the effect of relative humidity, mode, and the stage of the levitated droplet evaporation on bacterial survival and virulence. Overall, our study implied that the recipient exposed to the bacteria-laden droplet at high RH and early stage of the aerosol mode (contact free) of droplet evaporation is more prone to infection.

## Acknowledgements

Saptarshi Basu acknowledges financial support from the Science and Engineering Research Board (SERB), Pratt & Whitney Chair Professorship and SERB-SUPRA (Scientific and Useful Profound Research Advancement) project number SERB/F/10572/2021-2022. Dipshikha Chakravortty acknowledges financial support from DAE SRC fellowship (DAE00195), DBT-IISc partnership umbrella program for advanced research in biological sciences and Bioengineering. Dipshikha Chakravortty acknowledges infrastructure support from ICMR (Centre for Advanced Study in Molecular Medicine), DST (FIST), and UGC (special assistance) along with ASTRA-Chair fellowship, TATA Innovation grant, and DBT-IOE partnership grant. Amey Nitin Agharkar gratefully acknowledges the PMRF scheme facilitated by the MHRD, Government of India. Dipasree Hajra sincerely acknowledges the CSIR-SPM fellowship for her financial support. The JEOL-SEM IT 300 facility at AFMM, IISc and the MCB Confocal Imaging Facility, IISc. The funders had no role in study design, data collection, and analysis, publication decisions, or manuscript preparation.

## Author Contribution

Amey Nitin Agharkar – Conceptualization, Data curation, Formal Analysis, Investigation, Methodology, Validation, Visualization, Manuscript writing, reviewing and editing. Dipasree Hajra -Conceptualization, Data curation, Formal Analysis, Investigation, Methodology, Validation, Visualization, Manuscript writing, reviewing and editing. Kush Kumar Dewangan -Data curation, Formal Analysis, Investigation, Methodology, Validation, Visualization, Manuscript review and editing. Durbar Roy -Investigation, Methodology, Visualization, Manuscript review and editing. Dipshikha Chakravortty – Funding acquisition, Project Administration, Resources, Supervision, Manuscript review and editing. Saptarshi Basu-Funding acquisition, Project Administration, Resources, Supervision, Manuscript review and editing.

## Data availability

Data will be made available on request.

