## Supplementary Material for "Evaporation and pathogenesis of levitated bacteria-laden surrogate respiratory fluid droplets: At different relative humidity and evaporation stages"

### Supplementary Data

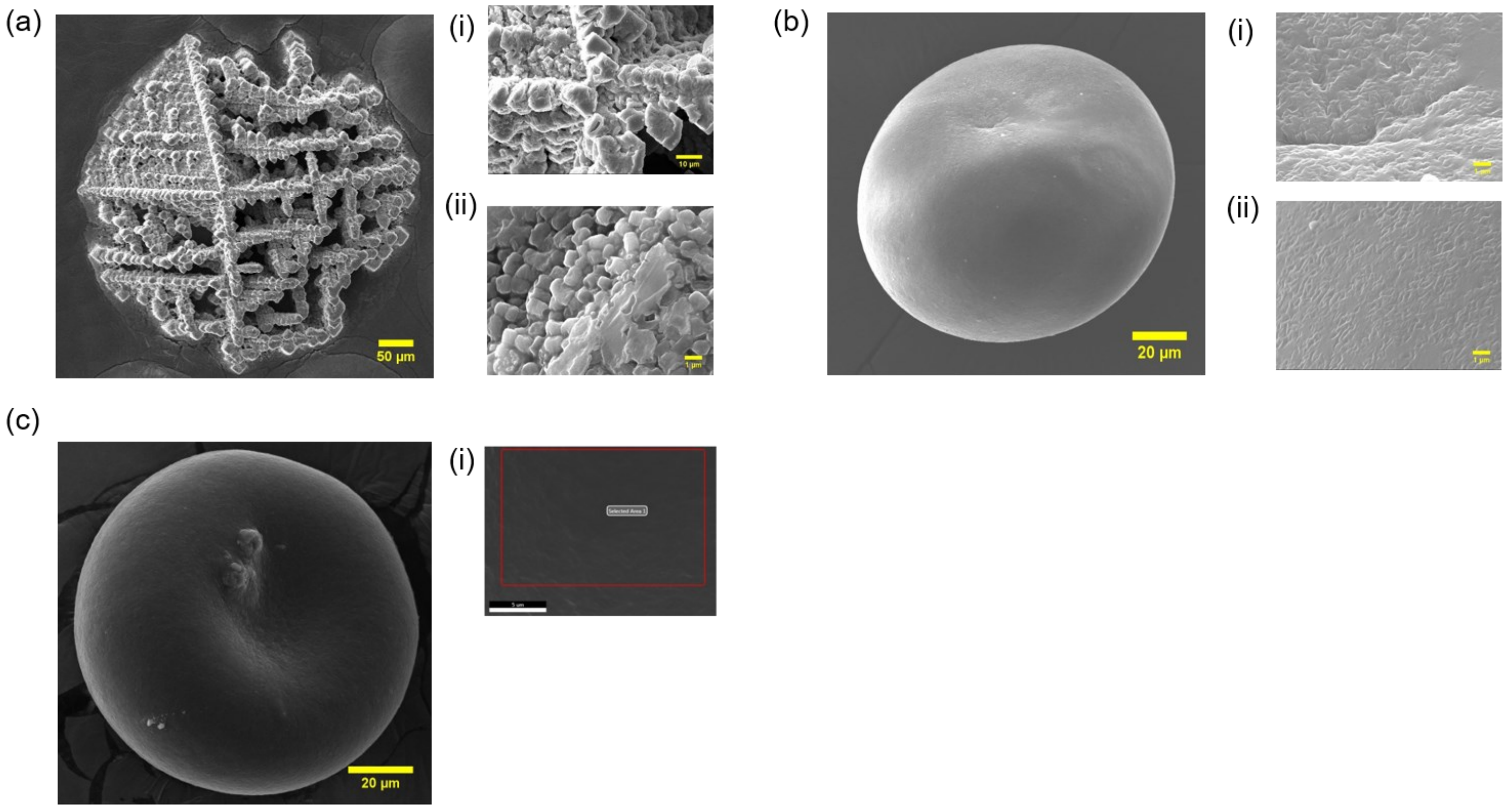


**Figure S1: SEM images of aerosol precipitates of (a) Low RH KP + 0.9% wt. NaCl (i) centre and (ii) edge (b) Low RH KP + 0.3% wt. Mucin (i) centre and (ii) edge (c) Low RH KP (i) centre**

(a)

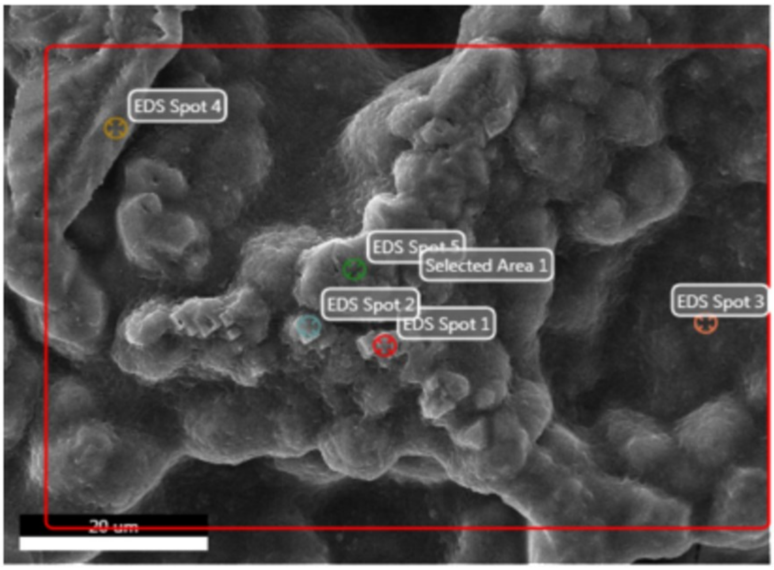


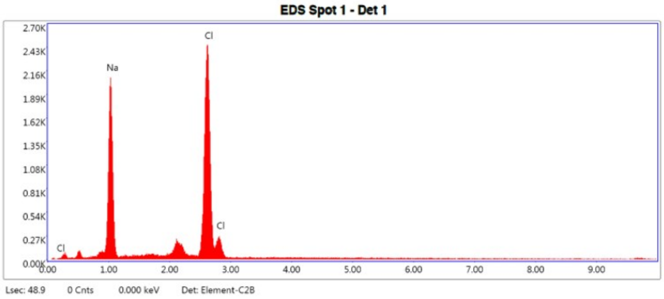


**eZAF Smart Quant Results**

| Element | Weight % | Atomic % | Net Int. | Error % | Kratio | Z | A | F |
|---|---|---|---|---|---|---|---|---|
| NaK | 36.19 | 46.65 | 338.13 | 5.36 | 0.2616 | 1.0478 | 0.6889 | 1.0015 |
| ClK | 63.81 | 53.35 | 550.88 | 2.62 | 0.5859 | 0.9719 | 0.9440 | 1.0007 |

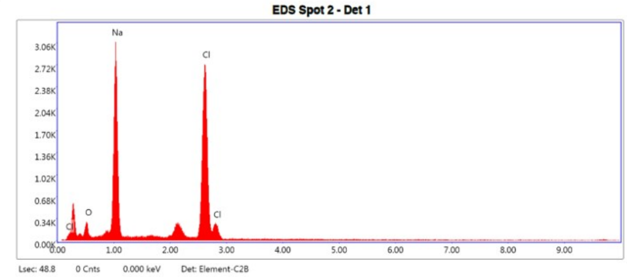


**eZAF Smart Quant Results**

| Element | Weight % | Atomic % | Net Int. | Error % | Kratio | Z | A | F |
|---|---|---|---|---|---|---|---|---|
| O K | 7.41 | 12.56 | 37.45 | 13.37 | 0.0191 | 1.1385 | 0.2258 | 1.0000 |
| NaK | 40.14 | 47.34 | 499.55 | 5.24 | 0.2815 | 1.0303 | 0.6796 | 1.0014 |
| ClK | 52.44 | 40.10 | 605.98 | 2.69 | 0.4695 | 0.9551 | 0.9361 | 1.0011 |

(b)

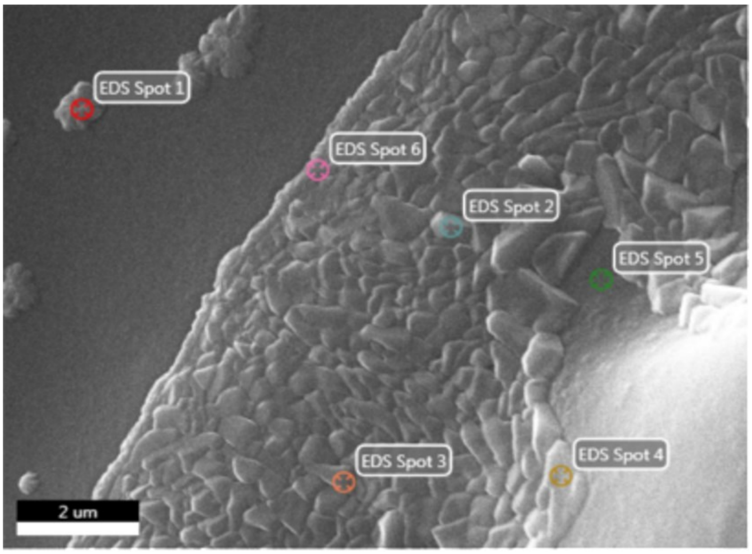


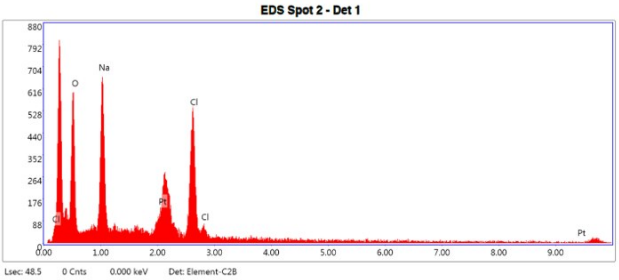


**eZAF Smart Quant Results**

| Element | Weight % | Atomic % | Net Int. | Error % | Kratio | Z | A | F |
|---|---|---|---|---|---|---|---|---|
| O K | 36.10 | 49.29 | 96.90 | 10.31 | 0.1150 | 1.0864 | 0.3521 | 1.0000 |
| NaK | 33.94 | 32.25 | 122.86 | 7.41 | 0.1614 | 0.9821 | 0.5810 | 1.0011 |
| ClK | 29.96 | 18.46 | 117.99 | 4.28 | 0.2132 | 0.9092 | 0.9377 | 1.0024 |

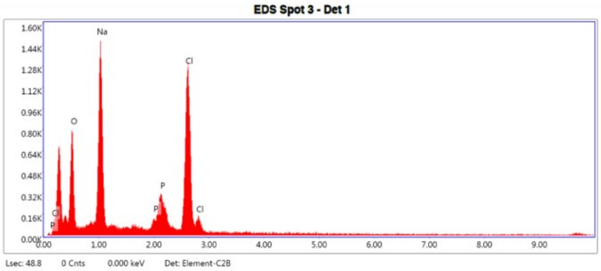


**eZAF Smart Quant Results**

| Element | Weight % | Atomic % | Net Int. | Error % | Kratio | Z | A | F |
|---|---|---|---|---|---|---|---|---|
| O K | 26.58 | 39.04 | 121.53 | 9.99 | 0.0864 | 1.1040 | 0.2943 | 1.0000 |
| NaK | 33.25 | 33.98 | 256.92 | 6.34 | 0.2022 | 0.9984 | 0.6084 | 1.0013 |
| P K | 3.84 | 2.91 | 34.74 | 5.15 | 0.0314 | 0.9554 | 0.8434 | 1.0134 |
| ClK | 36.33 | 24.07 | 288.20 | 3.25 | 0.3119 | 0.9248 | 0.9270 | 1.0018 |

**Figure S2: Energy-dispersive spectroscopy (EDS) report of aerosol precipitates of (a) Low RH KP + SRF Centre (i) Spot1 (ii) Spot 2 (b) Low RH KP + SRF Edge (i) Spot 2 (ii) Spot 3**

(a)

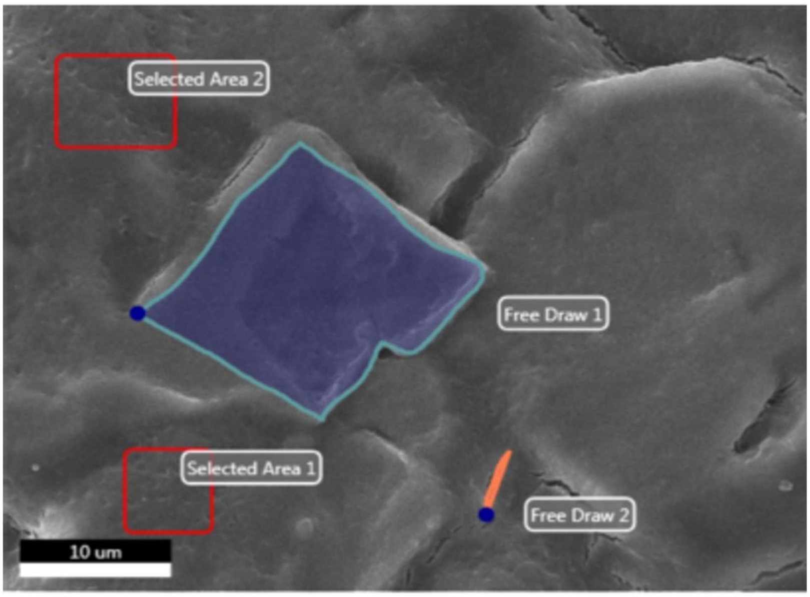


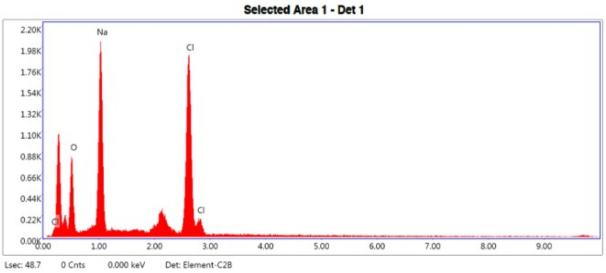


eZAF Smart Quant Results

| Element | Weight % | Atomic % | Net Int. | Error % | Kratio | Z | A | F |
|---|---|---|---|---|---|---|---|---|
| O K | 23.04 | 34.68 | 122.73 | 10.17 | 0.0705 | 1.1104 | 0.2753 | 1.0000 |
| NaK | 35.48 | 37.16 | 347.19 | 6.09 | 0.2203 | 1.0043 | 0.6173 | 1.0013 |
| ClK | 41.47 | 28.16 | 416.22 | 2.89 | 0.3627 | 0.9304 | 0.9384 | 1.0016 |

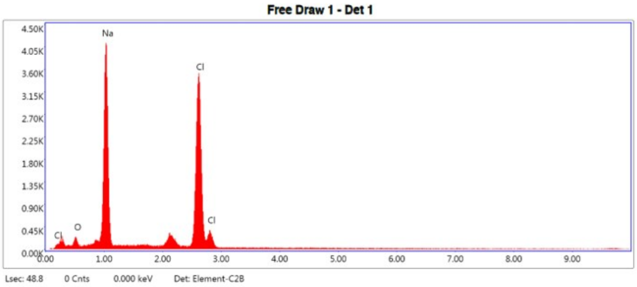


eZAF Smart Quant Results

| Element | Weight % | Atomic % | Net Int. | Error % | Kratio | Z | A | F |
|---|---|---|---|---|---|---|---|---|
| O K | 3.65 | 6.31 | 22.95 | 16.74 | 0.0093 | 1.1436 | 0.2218 | 1.0000 |
| NaK | 43.61 | 52.51 | 712.41 | 4.83 | 0.3176 | 1.0349 | 0.7028 | 1.0013 |
| ClK | 52.74 | 41.18 | 770.69 | 2.66 | 0.4719 | 0.9595 | 0.9316 | 1.0011 |

(b)

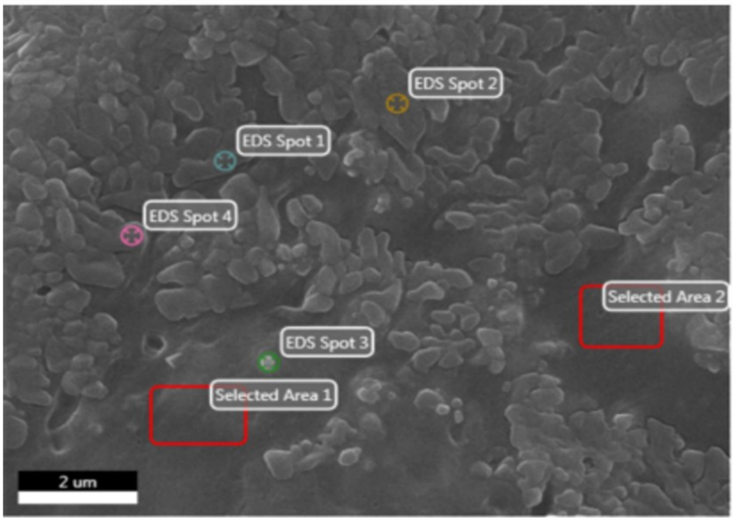


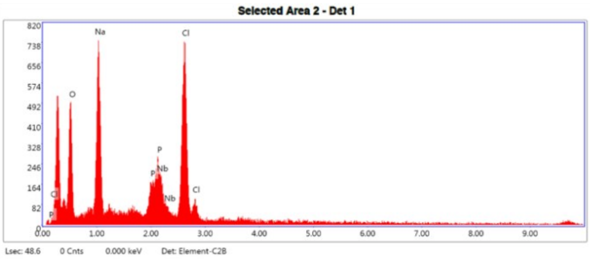


eZAF Smart Quant Results

| Element | Weight % | Atomic % | Net Int. | Error % | Kratio | Z | A | F |
|---|---|---|---|---|---|---|---|---|
| O K | 28.54 | 41.72 | 72.70 | 11.00 | 0.0740 | 1.1022 | 0.2917 | 1.0000 |
| NaK | 29.39 | 29.90 | 124.85 | 7.22 | 0.1405 | 0.9967 | 0.5939 | 1.0014 |
| P K | 6.64 | 5.02 | 34.20 | 9.58 | 0.0441 | 0.9539 | 0.8528 | 1.0128 |
| ClK | 35.43 | 23.37 | 157.45 | 3.85 | 0.2434 | 0.9233 | 0.9212 | 1.0018 |

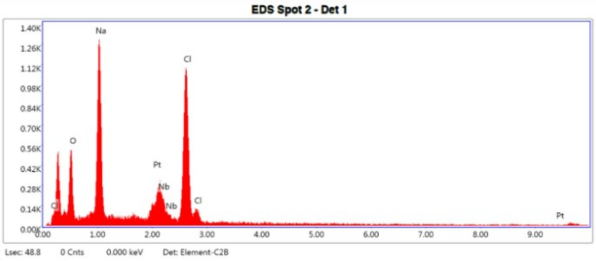


eZAF Smart Quant Results

| Element | Weight % | Atomic % | Net Int. | Error % | Kratio | Z | A | F |
|---|---|---|---|---|---|---|---|---|
| O K | 22.81 | 34.12 | 77.19 | 10.89 | 0.0578 | 1.1092 | 0.2844 | 1.0000 |
| NaK | 37.69 | 39.23 | 229.67 | 6.20 | 0.1900 | 1.0032 | 0.6254 | 1.0012 |
| ClK | 39.50 | 26.66 | 242.78 | 3.22 | 0.2759 | 0.9293 | 0.9353 | 1.0017 |

**Figure S3: EDS report of aerosol precipitates of (a) High RH KP + SRF Centre (i) Selected Area 1 (ii) Free Draw 1 (b) High RH KP + SRF Edge (i) Selected Area 2 (ii) Spot 2**

#### Section S1: p values and Statistical analysis

The statistical decision consists of accepting or rejecting the null hypothesis $H_0$ or the hypothesis of no difference [1]. This null hypothesis testing helps researchers to ascertain any conclusions regarding a population by examining sample from a population. It is rejected if the computed test value, in our case t test value, falls in the rejection region, and it is accepted if the computed value of the t test statistic falls in the non-rejection region. If it is rejected, we conclude that our alternate hypothesis $H_A$ (significant difference between two groups of variables) or our main research hypothesis is true. If our null hypothesis is not rejected, we conclude that any of the differences arising between the two cohorts of variables are not significant and are due to sampling error. The p value is a number that tells us whether our results are true based on the level of significance. The level of significance, α, is the probability of rejecting a true null hypothesis. A p value provides an indication about the reliability of the results.

test statistic = (relevant statistic - hypothesized parameter) /

standard error of the relevant statistic

To compute our test statistic t test, where $\bar{x}$ is the relevant statistic in our case mean of the variables and its standard deviation is $u_{\bar{x}}$ or the hypothesized parameter and $\sigma \,/\, \sqrt{n}$ is the standard error of the relevant statistic, we arrive at the following formula for transforming the normal distribution of mean to the standard normal distribution:

$$t = (\bar{x} - u_{\bar{x}})/(\sigma/\sqrt{n})$$

If the computed value falls into the rejection region, we accept the alternative hypothesis that is our results have significant differences or changes. The computed test statistic is represented with a p value. For example, if $p=0.05$ and the computed t test statistic value is 2.1, this denotes that the probability of getting a value greater than or lower than 2.1 is 0.05 when the null hypothesis is true. Therefore, when the p value is less than 0.05, null hypothesis is rejected, and we accept our research or alternate hypothesis. In our study, $p<0.05$ were considered significant.

$p<0.05$ *, $p<0.01$**, $p<0.001$***
